# Structural rationale to understand the effect of disease-associated mutations on Myotubularin

**DOI:** 10.1101/2022.07.27.501705

**Authors:** Teerna Bhattacharyya, Avishek Ghosh, Shailya Verma, Padinjat Raghu, Ramanathan Sowdhamini

**Affiliations:** National Centre for Biological Sciences, Tata Institute for Fundamental Research, Bangalore, India; Systems Biology Ireland, University College Dublin, Dublin, Ireland; Boston Children’s Hospital, Boston, MA, USA

## Abstract

Myotubularin or MTM1 is a lipid phosphatase that regulates vesicular trafficking in the cell. The *MTM1* gene is mutated in a severe form of muscular disease, X-linked myotubular myopathy or XLMTM, affecting 1 in 50,000 newborn males worldwide. There have been several studies on the disease pathology of XLMTM, but the structural effects of missense mutations of MTM1 are underexplored due to the unavailability of a crystal structure. MTM1 consists of three domains- a lipid-binding N-terminal GRAM domain, the phosphatase domain and a coiled-coil domain which aids dimerization of Myotubularin homologs. While most mutations reported to date map to the phosphatase domain of MTM1, the other two domains on the sequence are also frequently mutated in XLMTM. To understand the overall structural and functional effects of missense mutations on MTM1, we curated several missense mutations and performed *in silico* and *in vitro* studies. Apart from significantly impaired binding to substrate, abrogation of phosphatase activity was observed for a few mutants. Possible long-range effects of mutations from non-catalytic domains on phosphatase activity were observed as well. Coiled-coil domain mutants have been characterised here for the first time in XLMTM literature.

**Author Summary:** X-linked myotubular myopathy is a rare paediatric disorder and affected males suffer neonatal death or may live on only with ventilatory support. In this study, we employed a range of approaches to understand the molecular level effects of patient-derived mutations on an enzyme directly linked to the congenital muscular disorder. Using three-dimensional modelling and simulations, the effect of these mutations on the structure of the enzyme and its ability to bind its substrate was studied. To complement theoretical observations, experiments were performed with cells expressing this enzyme and its mutants. These studies reveal that each part of the protein may directly or indirectly affect its enzyme activity and most of the patient-derived mutations are expressed insufficiently in the cell. With the advent of genome sequencing technology, identification of congenital mutations is easier; computational studies of molecular consequences of mutations on a protein function such as this will prove immensely useful in understanding the disease and its prognosis.

## Introduction

Myotubularins are a large family of lipid phosphatases consisting of 14 paralogs, the first one to be identified being Myotubularin or MTM1. All the members have a GRAM domain (domain in Glucosyltransferases, Rab-like GTPase activators and Myotubularins), the lipid-binding domain and a phosphatase (PTP) domain. Although the myotubularins have a very low affinity for phosphotyrosine substrates *in vitro*, they share the overall structure of tyrosine phosphatases and the conserved HCX_5_R catalytic motif [1,2]. The cysteine acts as a nucleophile and dephosphorylates phosphoinositides at the 3-phosphate position. A coiled-coil domain is the additional domain present in 13 out of the 14 paralogs and implicated in aiding either homo or hetero-dimerisation of the myotubularins [3,4].

Myotubularin or MTM1 is a 603 amino acid long protein that possesses an N-terminal GRAM domain, followed by the catalytic phosphatase or PTP domain, a coiled-coiled domain and a C-terminal PDZ-binding motif, which also contains a PEST region [5]. Two important regions that lie within the catalytic domain are the Rac1-induced localisation to membrane ruffles (RID) and the SET-interacting domain (SID). While the former aids MTM1 to localise to membrane ruffles, the latter protein-protein interaction region is said to enable interaction of MTM1 and its homolog, MTMR12 [6]. MTM1 acts on the lipid substrates PI3P and PI(3,5)P_2_, which form small but distinct pools of lipids in endomembrane and are known to be key regulators of autophagy and membrane trafficking at the endosomal level [7–9]. PI5P is the product of dephosphorylation of PI(3,5)P_2_ and is involved in processes like oxidative stress signalling, growth factor signalling and transcriptional regulation [10]. In skeletal muscle cells, Myotubularin regulates the architecture of an intermediary filament called desmin, which maintains mitochondrial homeostasis through scaffolding [11].

Myotubular myopathy is a muscular disease condition first described in 1966 in an adolescent boy whose muscle biopsy revealed small myofibres and centrally placed nuclei, reminiscent of the foetal myotube stage of muscle development [12]. Initial analyses focused only on histopathological studies of affected males, but, the *MTM1* gene was first identified as mutated in X-linked myotubular myopathy (XLMTM) [5]. XLMTM is characterised by severe hypotonia and general muscle weakness, leading to respiratory disorders. It is primarily known to affect newborn males, and most XLMTM patients die at an average age of 4–8 months due to respiratory failure. The patients who may live depend entirely or partially on ventilator support [4,5,13,14]. With the advent of clinical genetics and improved exome sequencing data, several mutations from XLMTM have been identified, which map onto *MTM1*, and a few have been characterised in terms of lipid phosphatase activity, protein-protein interactions and effects on cell endosomal-lysosomal pathway [9,15–17]. Mutations associated with XLMTM have been found to map all over the MTM1 sequence, but most reported are from the GRAM and phosphatase domains [18]. Although the disease is fatal in most males in the neonatal or postnatal period, females carrying mutations in MTM1 were thought to be asymptomatic earlier. However, in the last few years, there have been a few studies on female carriers of XLMTM mutation, and late-onset symptoms such as the gradual decline of respiratory function have been observed, raising the possibility of X-linked haploinsufficiency in females [18,19].

In this study, a list of XLMTM mutations from patients with varying levels of disease severity [18,20–27] were curated for *in silico* and *in vitro* characterisation. For structural analyses, two models were created-one for the GRAM and phosphatase domains and a dimeric model of the coiled-coil domain. The structural and functional effects were assessed by molecular docking of the substrate (phosphoinositide 3-phosphate or PI3P) and interacting partners (desmin) to wild-type and mutants and by testing the protein expression and a LC-MS/MS based lipid 3-phosphatase activity *in vitro*.

## Results

### Homology models of MTM1 domains and selection of mutants spanning the entire protein sequence

The full-length structure of MTM1 or homologs within the Myotubularin lipid phosphatase family has not yet been reported. However, the crystal structure of MTMR2 (1ZSQ [3]) includes the GRAM and phosphatase catalytic domains and the aligned sequence stretches of MTMR2 and MTM1, corresponding to the GRAM and catalytic domains in both proteins, share close to 70% sequence identity. Using 1ZSQ as the template, a homology model of the MTM1 GRAM and catalytic domains was created (**Supplementary Figures 1A, B**), and the Ramachandran map and ProSA statistics were found to be in acceptable ranges (**Supplementary Figure 2**).

A coiled-coil region consists of two amphipathic helices and is formed from a repeat of the residue heptad pattern ‘abcdefg’, where ‘a’ and ‘d’ are conserved hydrophobic positions that form the dimerisation interface between two proteins, and e and g are polar residues which engage in electrostatic interactions. In the family of Myotubularins, coiled-coil is one of the domains present in all the members, irrespective of their catalytic activity status, and it helps the homologs form hetero- and homodimers. A dimeric model of the coiled-coil region of MTM1 was created using a multi-chain modelling approach.

A set of 140 XLMTM-associated mutations were curated from the literature and various resources on XLMTM patients and SNP records of *MTM1*(**Supplementary Table 1**). The effect of these single residue mutations on the function of MTM1 was assessed using SIFT and Polyphen-2, and out of the mutations that were found to “affect protein function” or to be “probably/possibly damaging”, a total of 13 were selected based on patient data, residue conservation, location on the MTM1 sequence. These 13 mutations were classified into four groups, corresponding to their phenotype, evolutionary conservation, frequency of mutation and structural and functional relevance (classes A, B, C and D, respectively) (**Figure 1, Table1**).

**Figure 1.**
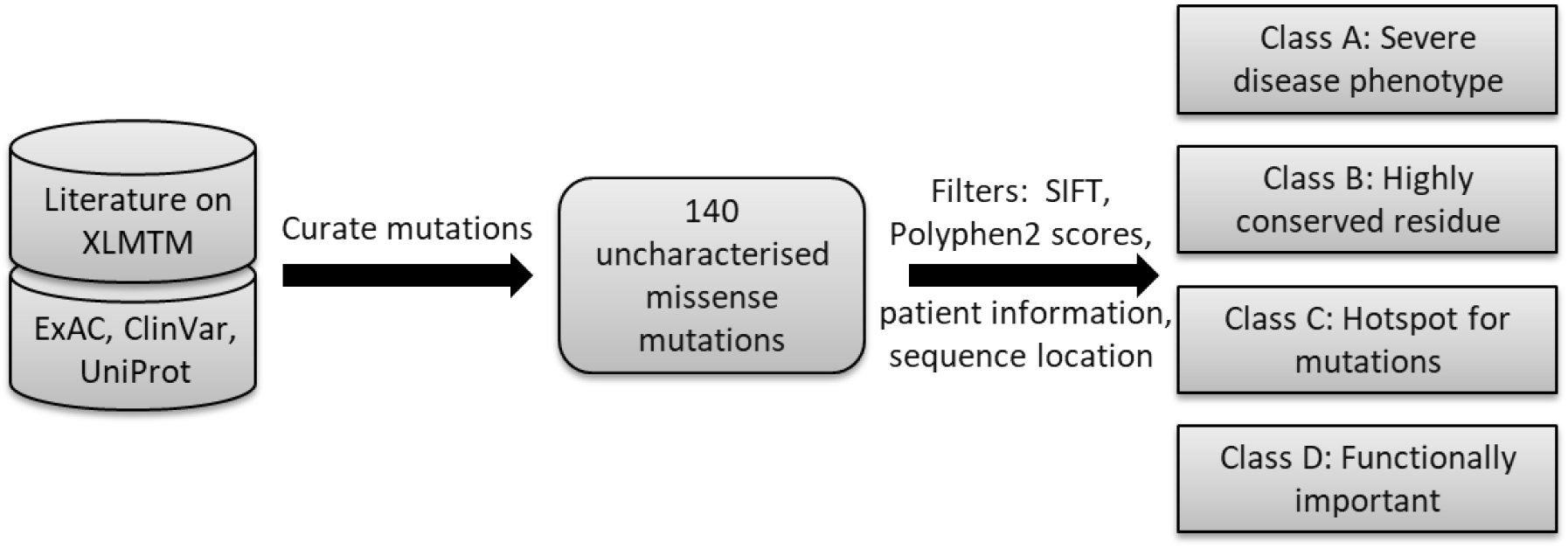
Selection of mutants implicated in XLMTM or predicted to be so. Previously uncharacterised missense mutations on the MTM1 gene were curated from literature and databases recording mutations on the *MTM1* gene, and these were grouped into four classes. For further analysis, three mutations were considered from classes A-C and four mutations from class D.

**Table 1.** List of mutations derived from patient-data and SNP databases mapping onto all the domains of MTM1 categorised according to the disease phenotype (A), evolutionary information and in silico predictions (B), hotspots for the mutation (C) and structural and functional implications (D). Several mutants have been obtained from recent papers (2003 and later) reporting X-linked Myotubular Myopathy patient data. The domain region indicates which domain/region of MTM1 the mutation falls on. The Polyphen-2 and SIFT scores are *in silico* measures of the effect of the mutation on MTM1 function- the closer the Polyphen-2 score is to 1, the more destabilising the mutation is, while the same applies for lower SIFT scores (closer to 0). Patient information reports the patient’s health status from whom the mutation has been derived. Residue conservation information was obtained by aligning vertebrate MTM1 sequences. The destabilisation of structure is indicated by the change of free energy of the protein (ΔΔG), and this was measured using the FoldX suite; More positive values of FoldX ΔΔG indicate destabilisation.

| **Mutation** | **Source** | **Domain**<br>**Region** | **Polyph**<br>**n-2 score** | **SIFT**<br>**score** | **Disease**<br>**severity** | **Residue**<br>**Conserv**<br>**ation** | **FoldXΔ**<br>**ΔG**<br>**(kcal/**<br>**mol)** |
| --- | --- | --- | --- | --- | --- | --- | --- |
| **Class A: Mutations with documented severe disease phenotype** |  |  |  |  |  |  |  |
| **C444Y** | Tanner<br>1999 [27] | Catalytic<br>Domain,<br>SID | 1 | 0 | Severe | Across all<br>vertebrates | 12.2524 |
| **I264S** | Buj-bello<br>1999 [22] | Catalytic<br>Domain | 1 | 0 | Severe | Across all<br>vertebrates | 3.61469 |
| **A389D** | Biancalana<br>2003 [21] | Catalytic<br>Domain | 1 | 0 | Severe | Across all<br>vertebrates | 7.64809 |
| **Class B: Mutations predicted to be damaging by several bioinformatics<sup>1</sup> tools** |  |  |  |  |  |  |  |
| **R220T** | Penon 2018<br>[18] | Catalytic<br>Domain | 1 | 0 | Maternal<br>mutation | Across all<br>vertebrates | 5.65505 |
| **W230C** | Tanner<br>1999,<br>Biancalana<br>2003<br>[21,27] | Catalytic<br>Domain | 1 | 0 | Severe | Across all<br>vertebrates | 5.25905 |
| **I65N** | Annoussam<br>y 2019 [20] | GRAM | 0.996 | 0 | Intermediat<br>e | Conserved<br>hydrophobi<br>c residue | 4.5584 |
| **Class C: Mutations at residues which are hotspots for mutation in XLMTM** |  |  |  |  |  |  |  |
| **R69S** | Biancalana<br>2003 [21] | GRAM | 0.999 | 0.03 | Severe | In<br>mammals | 4.05952 |
| **G402R** | Herman<br>2002, Flex<br>2002,<br>Tanner<br>1999<br>[23,25,27] | Catalytic<br>Domain | 1 | 0 | Severe | Across all<br>vertebrates | 6.69603 |
| **W346S** | Laporte<br>2000,<br>Werlauff<br>2015<br>[28,29] | Catalytic<br>Domain | 1 | 0 | Mild | Across all<br>vertebrates | 6.42884 |
| **Class D: Mutations at residues structurally or functionally important for MTM1** |  |  |  |  |  |  |  |
| **G378E** | Hane 1999<br>[24] | Catalytic<br>Motif | 1 | 0 | Severe | Across all<br>vertebrates | 19.7445 |
| **L470P** | Biancalana<br>2003 [21] | Catalytic<br>Domain,<br>SID | 1 | 0.04 | Severe | Across all<br>vertebrates | 8.28682 |
| **I502K** | Annoussam<br>y 2019 [20] | Catalytic<br>Domain | 1 | 0 | Severe | Across all<br>vertebrates | 5.58879 |
| **R564H** | ExAC [26] | Coiled-coil | 0.999 | 0 | Not<br>available | Across all<br>vertebrates | 1.23586 |

An additional mutant included in the analysis was the catalytically dead mutant, MTM1 C375S, used as a negative control in *in vitro* studies. Although the mutation C375S renders the enzyme inactive, the low FoldX ΔΔG value of -0.65 kcal/mol indicated no significant structural destabilisation of this mutant. The positive control was the WT protein itself; in literature, there are no reported mutants that display activity better than that of WT MTM1. The free energy change (ΔΔG in kcal/mol) of the MTM1 models for the GRAM and catalytic domain upon single-residue mutations predicted using the FoldX suite was significant in all cases (> 3 kcal/mol). The destabilisation predicted by FoldX may not be a proxy for the biological scenario as it only considers the local changes to the protein structure due to a single residue mutation (FoldX mutation causes changes to only the side-chain and not to the backbone). Thus, the ΔΔG values obtained using FoldX for different disease-associated mutants may provide a glimpse of the effect on protein function and stability of the structure.

Most of the XLMTM mutations map onto the GRAM and catalytic domains of MTM1 (**Figures 2A, B)**. The mutation R564H is from the ExAC database which reports variants of a gene with low allele frequency but without any associated XLMTM disease information-unlike other mutants characterised in this study. This residue is conserved across all vertebrate Myotubularins and falls in the dimer interface of the MTM1 coiled-coil domain (**Figures 2C, D**). Energy measurements for the WT and R564H models showed that R564H destabilised the dimer by 3.31 kcal/mol, which was similar to known disease-associated mutants from the other two domains of MTM1 (**Tables 1 and 2**).

**Figure 2.**
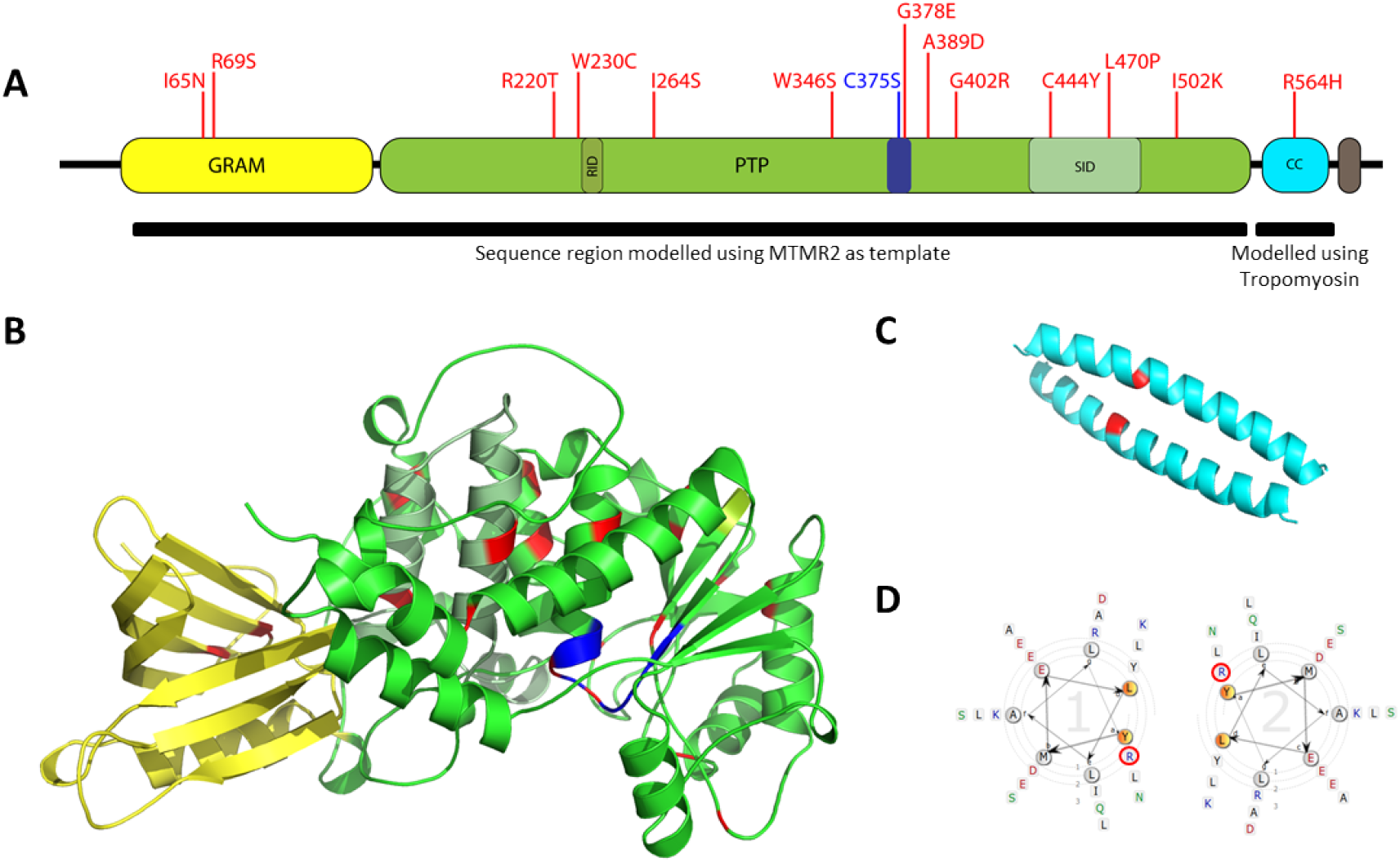
(A) The thirteen mutations (red) mapped onto the MTM1 sequence. A mutation in the catalytic motif-C375S (blue), which renders MTM1 catalytically inactive, was also included in the study as a control. MTM1 has an N-terminal GRAM domain (yellow), C-terminal coiled-coil domain (cyan) and a PDZ-binding motif (brown) apart from the catalytic domain (green). Within the catalytic domain, there are two regions-RID or Rac1 Induced recruitment Domain (dark green) and SID or SET-protein Interaction Domain (pale green) which are implicated in functions other than 3-phosphatase activity. (B) The mutations (red) have been mapped onto the model of the GRAM and PTP domains of MTM1. (C) Dimeric model of the coiled-coil region of MTM1, built using tropomyosin as a template; R564H mutation has been marked on the model. (D) Cartwheel representation of the sequences constituting the coiled-coil domain, indicating the R564H mutation (circled red) at the dimeric interface.

**Table 2.** COILCHECK+ is a tool designed for structural analyses of coiled-coil proteins [30,31]. The various energy parameters measured for the WT and mutant dimeric model of the coiled-coil domain of MTM1 show the destabilisation of the coiled-coil structure due to R564H mutation.

| **COILCHECK+**<br>**Energies** | **WT** | **R564H mutant** | **$\Delta E$ (kJ/mol)** | **$\Delta E$ (kcal/mol)** |
| --- | --- | --- | --- | --- |
| Hydrogen Bond Energy | 0 kJ/mol | 0 kJ/mol |  |  |
| Electrostatic Energy | -10.745 kJ/mol | -4.75 kJ/mol | 5.995 | 1.433 |
| van der Waals Energy | -292.35 kJ/mol | -284.487 kJ/mol | 7.863 | 1.879 |
| Total Stabilizing<br>Energy | -303.095 kJ/mol | -289.237 kJ/mol | 13.858 | 3.312 |

In this study, mutations within the catalytic domain that are not part of the catalytic motif but map to distant regions within the same domain have been included. Possible long-range effects of mutation (C444Y, A389D, R220T, G402R, L470P, W346S, L470P and I502K) on MTM1 structure and function were examined. The mutation G378E corresponds to mutation of a highly conserved glycine within the catalytic motif, and the mutations C444Y and L470P, which involve a change to a residue of similar chemical nature (hydrophobic), fall on the putative SET-interacting domain or SID, which is reportedly a protein-protein interaction domain of MTM1. Apart from the mutations in the catalytic domain and coiled-coil domain, two mutations (R69S, I65N) fall onto the putative PI5P binding region within the GRAM domain. Two mutations (W230C and I264S) map to the desmin-binding region of MTM1.

### Most mutants at the catalytic domain bind poorly to PI3P ligand

MTM1 has three main domains: the N-terminal GRAM, phosphatase or PTP, and C-terminal coiled-coil domains. The former two are involved in lipid binding and dephosphorylation, respectively. The catalytic motif of MTM1 is the same as the evolutionarily conserved motif of cysteine-based tyrosine phosphatases-HC(X)_5_R, where C is the catalytic cysteine (C375 in MTM1) and X is any amino acid. Although structurally similar to tyrosine phosphatases, MTM1 acts upon the relatively less abundant pool of phosphoinositide 3-phosphate (PI3P) and also phosphoinositide-3,5-bisphosphate (PI(3,5)P_2_), hydrolysing the 3-phosphate.

The homology model of the MTM1 GRAM and catalytic domain regions was generated using the crystal structure of MTMR2, a close homolog of MTM1, as the template. The structure of short acyl chain-containing PI3P or diC4 PI3P (bound to MTMR2 in the crystal structure [32]) was used as the ligand for docking with the MTM1 models (WT and 13 mutants from GRAM and catalytic domains) (**Figures 3A, B**). DiC4 PI3P is a phosphoinositide molecule with four-carbon acyl chains on the sn-1 and sn-2 positions, and a phosphodiester bond-linked inositol ring on the sn-3 position. There are two phosphate groups on diC4 PI3P, one on the phosphodiester link (D1-phosphate) and the other one on the 3^rd^ carbon position of the inositol ring (D3-phosphate) with the remaining positions containing hydroxyl groups (**Figure 3A**). A ligand-protein complex is stabilised by favourable non-covalent interactions between the ligand and residues within the protein’s binding pocket. These include hydrophobic interactions, aromatic interactions and hydrogen bonding interactions. The residues involved in the interaction with ligand in the WT and 11 mutant MTM1-docked complexes were traced to track changes in interacting partners owing to a mutation in the catalytic domain of MTM1 (**Figures 3A, D**). As seen from **Figure 3D**, the interacting partners and the nature of interactions change between the WT and mutant proteins. For example, the residue W379 (conserved across active myotubularin family members) forms a hydrogen bond with the 5’-OH of the ligand only in WT and mutant G378E but not in any other docked complexes. Similarly, the residue D380 from the catalytic motif forms one hydrogen bond each with the 5’ and 6’-OH in the WT and G378E docked complexes but fails to form more than one hydrogen bond in the mutants C375S, R220T, L470P and A389D and any interaction at all in rest of the mutants. The residue R381 housed within the catalytic motif and fully conserved across all myotubularins forms a bidentate hydrogen bond with the D1-phosphate and another hydrogen bond with the 6’-OH of the diC4 PI3P in the mutants C375S, R220T, L470P and A389D. On the other hand, some residues do not interact with the ligand in the WT MTM1-PI3P complex but engage in hydrogen bonding (Њ) or hydrophobic interactions (Φ) with the lipid in some of the mutants (R280, N284, A432, and R433). The highly conserved residues R280, N284, N288 and N313 (>60% conservation across MTM1 homologs), distant from the catalytic motif in sequence, do not interact with the D3-phosphate in WT MTM1. However, in several mutants, they engage in hydrogen bonding or hydrophobic interactions with the hydrolysable phosphate group. This could indicate stronger binding of the lipid substrate, leading to unsuccessful dephosphorylation by the disease-associated mutants. Overall, the traced interactions between substrate and receptor binding site (**Figure 3D**) highlight that binding is not limited to the catalytic motif residues; even the residues that are seemingly distant from the motif also play important roles in substrate binding, hydrolysis and release.

**Figure 3.**
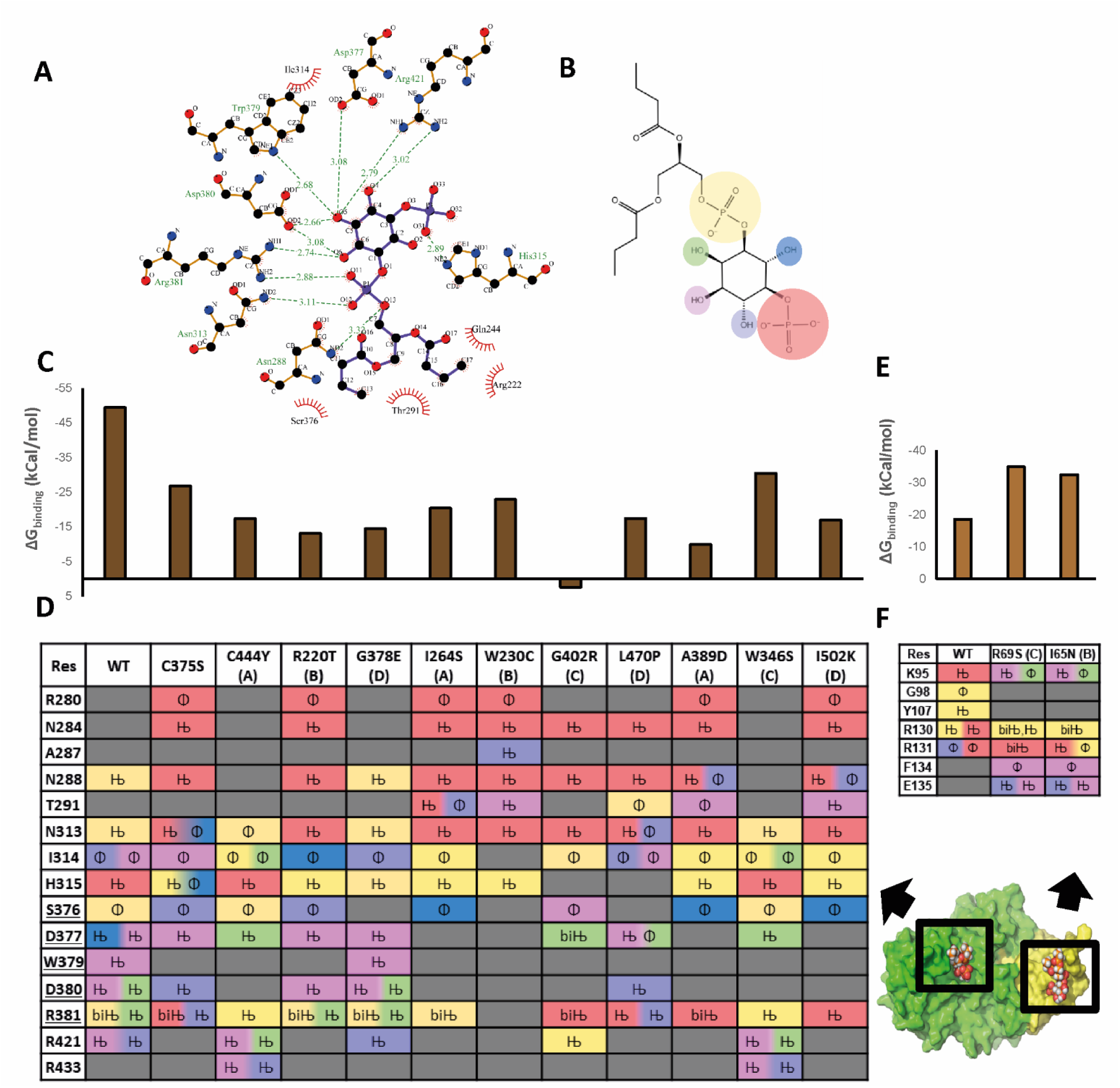
(A) Protein-ligand interactions visualised in 2D, showing hydrogen bonds (green dashed line) and hydrophobic contacts (brown circle) between PI3P (purple backbone) and MTM1 residues (yellow backbone). (B) The chemical groups of the ligand (PI3P) are colour-coded to indicate contacts with MTM1 residues at the binding site. (C) ΔG_binding_ (kcal/mol) for WT and mutant MTM1-PI3P complexes (at the catalytic site). (D) Residues interacting with PI3P, nature of the interaction (HBond: Њ and Hydrophobic: Φ) and a chemical group of ligand involved in the interaction at the docking site within the PTP domain. One residue may interact with multiple chemical groups in different modes of interaction. Residues underlined are those which are part of the catalytic motif of PTP. (E) ΔG_binding_ (kcal/mol) for WT and mutant MTM1 and PI3P complexes (with the GRAM domain). (F) Interactions between GRAM domain residues and ligand seen for WT and mutant complexes. (G) Space-filling model showing the GRAM domain (yellow) and PTP domain (green) binding sites to which PI3P was docked and the docked poses were analysed (D, F).

Trends in the predicted docking score and free energy of binding or ΔG_binding_ values of the MTM1-PI3P docked complexes (**Supplementary Table 2**) provide valuable insight into the stability of the mutant receptor complexes compared to WT MTM1. For almost all mutant complexes, the docking score was found lesser than the WT MTM1-PI3P complex. It was easily discernible from ΔG_binding_ values that WT MTM1 forms the most stable complex with the substrate, PI3P, at its catalytic domain. The catalytically inactivating C375S mutation reduces the ΔG_binding_ to about half of that WT. A similar reduction is seen for almost all other mutants, including sites away from the catalytic motif (**Figure 3C, Supplementary Table 2**). Interestingly, the binding site of the R220T docked complex resembled C375S closely, although the mutation is not from the catalytic motif (**Figure 3D**).

Additionally, the hydrophobic contacts between the carbons of the inositol ring of diC4 PI3P and the different residues within the binding pocket in the PTP domain were traced across the mutants. In several mutants, the residues R381 and D433 interact with the cyclohexane ring but not in the WT docked complex (**Supplementary Figure 3**). Similarly, the mutants gather more interactions with the inositol ring (residues I314 and R421) compared to the WT. This alludes to stronger binding of the lipid substrate, leading to unsuccessful dephosphorylation.

Apart from the catalytic domain, which hydrolyses the D3-phosphate of PI3P, the GRAM domain also possesses an affinity for PI3P and is believed to guide the protein to distinct subcellular pools of PI3P. The precise binding site of PI3P in the GRAM domain of MTM1 has not been discussed earlier, but the role of two loops in the GRAM domain in interaction with PI3P had been alluded to in the structural analysis of MTMR2 [33]. To validate this extrapolation from MTMR2 to its close homolog, MTM1, blind docking of the PI3P ligand with the MTM1 model was carried out. In this exercise, the ligand was free to interact with any region of the protein. In several poses obtained, the ligand was bound to the GRAM domain through interactions with residues from the same two loops connecting the β5-β6 and β7-α1 within the GRAM domain (**Supplementary Figures 4A, B**). Through careful analysis of the surface electrostatic charge distribution on the MTM1 model, it was observed that the region is rich in positively charged residues and lies on the same face as the active site (**Supplementary Figures 4C**)-we propose that this is the putative PI3P binding site within the GRAM domain.

### MTM1 mutations away from catalytic site affect the ability to bind cognate lipid and protein substrates in different ways

Next, the same diC4 PI3P ligand was docked to the GRAM domain, defining the grid using residues identified to interact with the ligand through blind docking. The interacting residues were traced for the WT MTM1-PI3P docked complex and the two GRAM domain mutants. It was interesting to note that the GRAM domain mutants R69S (corresponds to a hotspot mutant, class-C) and I65N (class-B) engage in additional interactions with the ligand compared to WT (**Figure 3E**) and form a more stable complex as shown by lower ΔG_binding_ values (more negative values) (**Figure 3F**). It is speculative whether the more favourable binding of PI3P to the mutant proteins may lead to the inability of the GRAM domain to “pass on” the lipid substrate to the catalytic domain, decreasing the efficiency of dephosphorylation, compared to that of WT MTM1.

Apart from the phosphatase activity, MTM1 binds to intermediary filament desmin in skeletal muscles and regulates the filament assembly shape and branching. The desmin-binding region of MTM1 is within the catalytic domain but away from the site of phosphatase activity (**Supplementary Figure 5A**). Two patient-derived mutants W230C and I264S map to this region of MTM1. We performed molecular dynamics simulations of the docked complexes of the WT and mutants to predict possible effects of these two mutations on the protein-protein interactions between MTM1 and desmin. **Supplementary Figure 5B** shows the residues from MTM1 and desmin chains engaged in several non-covalent interactions that stabilise the WT MTM1-desmin dimer complex. As seen in **Supplementary Figure 5C**, the Cα RMSD of the MTM1 chains remain relatively stable throughout the simulations (100ns) while the desmin chain Cα RMSD undergoes comparatively more significant fluctuations. While the Cα RMSD (Å) fluctuations of desmin stabilise beyond 60ns of the simulation for the WT MTM1-desmin complex, the Cα RMSD does not stabilise even till the end of the simulation in the case of W230C and I264S. When the number of hydrogen bonds formed between desmin and MTM1 was tracked over time, it was clear that WT MTM1 retains the highest number of hydrogen bonds (average 14) throughout the simulation than both of the MTM1 mutants (8 for W230C and 7 for I264S) (**Supplementary Figure 5D**). It was also seen that the mutant complexes show an increase in the radius of gyration (reflects compactness of the complex) over time (**Supplementary Figure 6A and B**). These observations point to the instability of the W230C and I264S MTM1-desmin complexes and impaired scaffolding activity.

### Biochemical measurements reveal the loss of 3-phosphatase activity in some MTM1 mutants

We performed site-directed mutagenesis on the FLAG-tagged MTM1 WT construct to obtain the mutant constructs. We also sub-cloned a shorter construct corresponding to the GRAM and catalytic domains of MTM1 into a FLAG-tagged vector, termed as MTM1ΔCC in this study. The protein expression was assayed by western blotting of the lysates from HEK293T cells transiently transfected by FLAG-tagged WT and mutant MTM1 plasmids. Varying levels of expression were observed for the WT and mutant proteins. While the mutants C375S, R220T, R69S, G378E and R564H expressed as much or even more than the WT MTM1 in the HEK293T cells, all other mutants showed poor expression (**Supplementary Figures 7 and 8**). To note from **Table 1**, that most of the mutants expressing poorly are observed to have a severe or damaging effect on the protein (as seen from the free energy change predicted by FoldX).

Function-test experiments were carried out to examine the phosphatase activity of the mutants. PI3P lipid 3-phosphatase activity was measured using a highly sensitive mass spectrometry-based assay for the mutants that expressed protein at least as much as the level of WT MTM1 using our transient transfection system. Cell lysates from mammalian cells transfected by respective plasmids were incubated with commercially available 37:4 PI3P, distinct from the endogenous pool of PI3P in mammalian cells, and subjected to phosphatase assay conditions (please see **Materials and Methods**). The reaction was performed for 15 minutes after which the lipids were extracted, derivatised and measured using AB Sciex 6500 triple quadrupole mass spectrometer detecting the lipids in MRM mode. The PI3P lipid 3-phosphatase activity is expressed in terms of “response ratio” which is the ratio of the amount of product signal (37:4 PI) to the amount of substrate signal (37:4 PI3P). While the untransfected control (UTC) lysates failed to elicit a decent response ratio, the WT MTM1 consistently produced response ratios around 0.5 (**Figure 4C**). In sharp contrast to this, quite expectedly, the catalytic dead C375S mutant lysates could not evoke a significantly different response compared to UTC (**Figure 4C**) [34].

**Figure 4.**
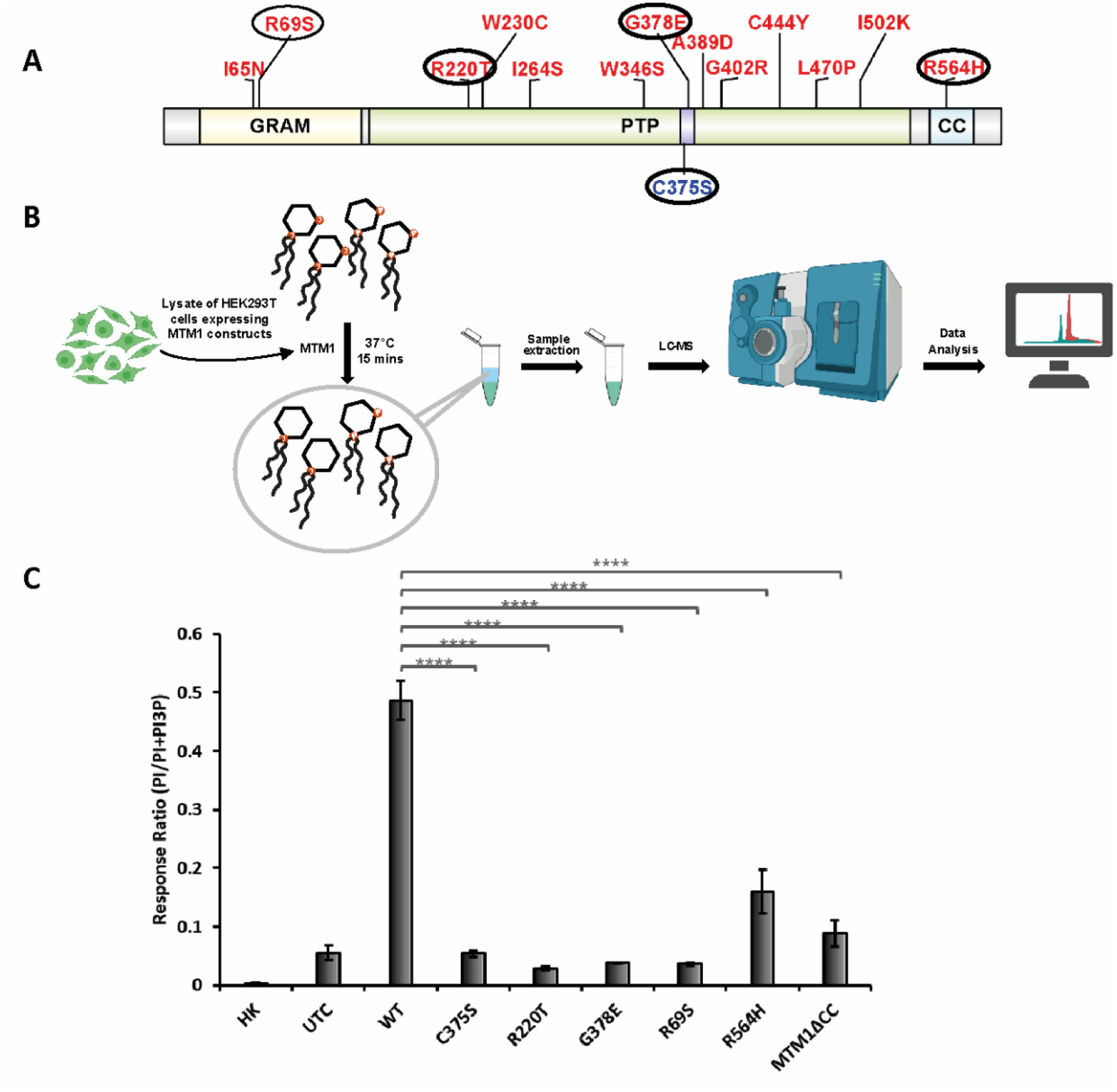
Abrogation of catalytic activity of MTM1 mutants towards PI3P substrate, measured using mass-spectrometry based assay (A) Mutants for which 3-phosphatase assay was carried out shown on the MTM1 sequence (circled in black). There is one mutant each, from classes B and C (R220T and R69S, respectively) and two mutants from class D (G378E and R564H). The class-A mutants expressed too poorly to be considered for activity measurement (B) Schematic representation of the phosphatase assay carried out for lysate of HEK293T cells expressing MTM1 WT or mutants (or shorter construct MTM1ΔCC, corresponding to only GRAM and catalytic domains) using 37:4 PI3P as substrate. The 3-phosphatase activity is expressed as the response ratio. (C) Activities of WT MTM1 and disease-associated mutants R220T, G378E, R69S and predicted mutant R564H and truncated construct MTM1ΔCC towards 37:4 PI3P. HK is a heat-killed WT MTM1 sample, and UTC corresponds to untransfected control. The mutants and the shorter construct show significantly lower 3-phosphatase activity than MTM WT towards PI3P substrate (**** indicates p-value < 0.0001 measured using one-way ANOVA with posthoc Tukey’s multiple comparison test).

R220T is a mutation from Class-B (Table 1) identified in a mother who suffered multiple neonatal losses and is a carrier for XLMTM [19]. Although R220T mutation is in a site distant from the catalytic motif and PI3P binding site, the mutant cannot form a stable docked complex with the ligand, as seen from docking studies (**Figures 3C and D**). This is reflected in the response ratio in the assay which shows complete inactivity towards PI3P lipid substrate (**Figure 4C**).

The mutation G378E (Class-D) was derived from a male newborn who suffered profound hypotonia and impaired respiratory effort and died at two months [24]. The residue G378 is part of the catalytic motif conserved among all Cys-based phosphatases, HCX_5_R, which in all active members of the Myotubularin family is VHCSDGWDRT. Substitution of the small hydrophobic residue with a bulkier charged residue is bound to have implications in the phosphatase activity of MTM1, and the same is reflected by the complete abrogation of catalytic activity of MTM1 G378E (**Figure 4C**).

R69S (Class-C) is a mutation derived from a male patient alive with 24 hours ventilator support at the time of detection of the mutation, but with severe XLMTM phenotype [9]. This positively charged residue has been implicated in putative PI5P binding of MTM1, leading to oligomerisation [18]. The initial attempt at measurement of the 3-phosphatase activity of R69S and two constructs associated with the coiled-coil region (R564H- a mutation at the dimeric interface and a shorter construct MTM1ΔCC) showed similar or higher activity than WT MTM1 (data not shown). In previous reports of MTM1 activity measurements, few mutants of MTM1 have been reported to express higher than WT, but activity reported has not been normalised to the protein expression. Quantitation of protein expression of the mutants revealed that while R220T and G783E express similar to WT MTM1, the mutants R69S and R564H and the MTM1ΔCC construct express much higher than WT (**Supplementary Figure 8**). The activity measurements were then performed after normalising the corresponding mutant expression to WT MTM1 expression. It was found that R69S, R564H and MTM1ΔCC are catalytically less efficient than WT (**Figure 4C**). This is, to our knowledge, the first time that the expression of mutants has been accounted for, for the measurement of MTM1 catalytic activity. Apart from the mutant G378E, which maps on to the catalytic motif itself, mutants at regions far from the catalytic motif (R69S, R220T and R564H) also displayed abrogation of phosphatase activity, and this goes to show the role of long-range effects residues or role of additional domains of MTM1 in its catalytic activity.

## Discussion

We selected a few mutations of MTM1 for this study, of which all but one are derived from X-linked myotubular myopathy disease. We employed multiple *in silico* approaches to quantify the destabilisation of MTM1 structure, substrate-binding and protein complex formation ability by these mutants, and these observations were augmented by *in vitro* experiments.

Patient-derived mutations mapping onto the GRAM and catalytic domains and an ExAC database derived mutation from the coiled-coil domain were curated for this study. To the best of our knowledge, any mutant from the coiled-coil region of MTM1 has not been characterised previously. To circumvent the absence of a crystal structure of MTM1, a homology model of the first two domains and a dimeric model of the coiled-coil domain were created, and the effect of mutations on the structure was measured as free energy change upon mutation. Docking PI3P to the catalytic domain of MTM1 revealed interesting observations-all mutants are unable to form stable complexes, some show stronger binding to the hydrolysable phosphate group of the substrate, alluding to inefficient dephosphorylation and long-range effects of mutations distant from the catalytic site. Through blind docking studies and inspection of the electrostatics of the MTM1 surface, a putative PI3P binding pocket within the GRAM domain on the same face as the catalytic domain was identified and is discussed here for the first time. In the case of the two mutants (I65N and R69S) mapping to the GRAM domain, a docked complex more stable than WT MTM1 was found-it may be speculated that tighter binding of the substrate by accessory domain ultimately affects the phosphatase activity by the catalytic domain. Additionally, molecular dynamics studies of desmin-MTM1 protein complexes showed impaired scaffolding activity in the case of two disease-associated mutants of MTM1.

Most of the disease-associated mutants (especially those from class-A) express poorly in mammalian cells, compared to WT MTM1 and could not be taken up for phosphatase activity measurement. The absence or low expression of MTM1 has been correlated to the XLMTM disease context, simply explained by the unavailability of the gene product in the cell [34,35]. The poor expression of several disease-associated mutants could be due to various reasons; misfolding of a protein or slight conformational changes due to a non-synonymous mutation directing it to proteasomal degradation. Missense mutations such as C444Y, A389D, G402R, W346S and L470P were predicted to cause a significant destabilisation of the MTM1 structure by 6-12 kcal/mol. This destabilisation is reflected in their poor expression in mammalian cells compared to WT. However, such correlation between predicted ΔΔG values and protein expression is not straightforward due to limitations of the prediction methods and other factors that lead to low expression, including misfolded mRNA structure and codon bias. The inability to form a stable complex with the cognate substrate may also render a cytosolic protein susceptible to proteasomal degradation, resulting in poor expression. Through docking studies, it was seen that several mutants are unable to form a stable docked complex (ΔG_binding_ of these mutants were significantly higher than the WT-PI3P docked complex), and they showed poor expression in transfected mammalian cells as well.

Lysates of mammalian cells expressing the different constructs of MTM1 were incubated with exogenous PI3P substrate to measure the 3-phosphatase activity using sensitive mass spectrometric techniques. Abrogation of 3-phosphatase activity was clearly observed for the mutants R69S, R220T, G378E, R564H and MTM1ΔCC, of which the latter two are changes associated with the coiled-coil domain of MTM1. It is noteworthy that R220T and G378E not only failed to form stable docked complexes with the substrate (PI3P) but are also catalytically inactive.

We infer from the above observations that the catalytic activity of MTM1 is affected by long-range interactions with residues that are not necessarily within the active site. The coiled-coil domain is known to be the dimerisation module of Myotubularin, but the implication of the loss of this domain or mutation destabilising this domain is being discussed here for the first time. To conclude from the results from the docking studies and biochemical assays- the accessory domains of MTM1 act synergistically with the catalytic domain for the phosphatase activity of this protein.

A majority of past studies on X-linked myotubular myopathy have looked into the effect of MTM1 mutations from a cell biological perspective. Possibly due to the unavailability of a full length MTM1 structure, there is a paucity of studies on the effects of mutations on MTM1 structure, linked to its function. We also noted a lack of *in silico* studies in the field of XLMTM and used structural bioinformatics approaches to study the effect of disease-associated mutations on a MTM1 structural model and augmented these with biochemical assays where appropriate. In the future, this pipeline can be used to screen disease-associated mutations and correlate them to the disease severity. Another interesting possibility to explore in future would be screening out mutants in which lipid phosphatase activity is not correlated to the disease. Although there has been significant advancement towards studying gene therapy as a treatment for XLMTM, there is a lacuna of understanding the exact link between MTM1 mutations and disease prognosis and severity, which we believe can be filled with such *in silico* approaches preceding *in vitro* or *in vivo* studies.

## Materials and Methods

### Selection of mutants and classification

A list of missense mutations of *MTM1* was compiled from ClinVar [36], UniProt [37] and ExAC [26] databases, as well as several publications reporting mutations found in X-linked myotubular myopathy patient samples. There have been a few biochemical characterisation studies of MTM1 mutants in the context of XLMTM. All mutations compiled were filtered to exclude such previously characterised mutants. A total of 140 single residue mutations were obtained, for which SIFT [38] and Polyphen-2 [39] scores were calculated; these are tools to predict the effect of an amino acid substitution on the sequence, structure and function of a protein. Single residue mutations annotated as “possibly/probably damaging” by Polyphen-2 and also “affects protein function” by SIFT were further filtered based on the availability of patient information (i.e., the severity of the XLMTM mutation in terms of patient health) and where the mutation mapped on to, on the MTM1 full-length sequence. These were classified into four main classes-(A) mutations which corresponded to severe disease phenotype, i.e. cases of early neonatal death or complete ventilator support upon survival, (B) degree of conservation of the residue across myotubularin family members, indicated by SIFT and Polyphen-2 scores, (C) mutations at hotspot residue locations, i.e. residues mutated frequently in XLMTM, and (D) mutations known to be or which could be structurally or functionally important for MTM1. The latter includes a mutation from the ExAC database, which has low allele frequency indicating a rare variant but does not have associated patient information, and it maps on to the coiled-coil domain- a region previously not much explored in terms of XLMTM disease. The conservation of the residues mutated in the list consolidated was checked by aligning MTM1 sequences from a few representative organisms (human, orangutan, mouse, rat, cow and *Xenopus laevis*) using CLUSTALO [40].

### Homology modelling of different domains of MTM1 and docking of phosphoinositide lipids

The GRAM domain of MTM1 is a lipid-binding domain and helps in the membrane targeting of MTM1 [17]. The phosphatase domain consists of the tyrosine phosphatase signature catalytic motif with a nucleophilic Cys. A crystal structure of the GRAM and catalytic domain of MTMR2 is available, with a short acyl chain containing PI3P bound to the active site of MTMR2 (PDB ID: 1ZSQ) [32]. Whereas MTM1 and MTMR2 full-length sequences share 65% sequence identity, the sequence identity for the aligned region, i.e. the GRAM and phosphatase domains, is a bit higher (close to 70%). 1ZSQ was used as a template for homology modelling of residues 32-543 of MTM1 using MODELLER [41]. Unless otherwise specified, this model of MTM1 excluding termini and the coiled-coil domain is referred henceforth as the MTM1 model.

A coiled-coil domain typically consists of heptad repeats of hydrophobic and hydrophilic residues. In previous work from the lab on the modelling of coiled-coil domains, heptad-to-heptad alignment of the sequence of the predicted coiled-coil domain was carried out with the sequence of a known coiled-coil structure [30]. The alignment was then used for modelling the three-dimensional structure of the coiled-coil domain of MTM1. Using Predicted coiled-coil and PAIRCOIL2 tools [42], the coiled-coil region was predicted to be within residues 551-584. With the predicted heptad information, an alignment was constructed with the sequence of Tropomyosin corresponding to the structure of C-terminal Tropomyosin fragment (PDB ID: 2EFR), and homology modelling was performed to obtain the dimeric coiled-coil segment of MTM1 using MODELLER [43].

Energy minimisation using the Impref module (Schrödinger suite) with a default cut-off RMSD value of 0.3 Å using the OPLS3e force field [44] [45] [45] [45] and assignment of side-chain tautomersat pH of 6.5 (experimentally determined pH optimum for MTM1 phosphatase activity) were carried for the MTM1 GRAM-PTP domain model as well as coiled-coil domain models. The quality of the models was assessed using the two tools-PROCHECK [45] generates a Ramachandran map of the residues of the model and ProSA [46] evaluates model quality using knowledge-based potentials of mean force, indicated by a Z-score. Stabilising energy for the coiled-coil dimer models were measured using the in-house COILCHECK+ program [30,31].

*In silico* mutagenesis was carried out using the FoldX suite (version 4) [47], and the change of ΔG in kcal/mol (ΔΔG) owing to each single residue mutationwas measured using in-house python scripts analysing FoldX results (Pankaj Chauhan Github: https://github.com/chauhan7892/Cardiac_Mutant). The RepairPDB function within FoldX was used to carry out energy minimisation of the mutated models using the empirical FoldX force field [47]. The MTM1 models (WT and mutants) thus generated were used as receptors for docking PI3P ligand.

The extra precision or XP glide docking method optimises the fit of the ligand to the receptor’s binding site (where the residue side-chains are rigid) in an induced manner while optimising computational time. For reference, the ligand obtained from the MTMR2 crystal structure (1ZSQ) was docked back to the MTMR2 structure (the receptor grid was defined as the residues within the catalytic motif) and several residues found interacting with the ligand in the top docked pose matched with those mentioned in the crystal structure [32]. Thus, XP glide docking was the method of choice for docking PI3P to MTM1 WT and mutant models [48]. The models were refined using the Impref energy minimisation module within the Schrödinger suite with a default cut-off of 0.3 Å RMSD using the OPLS3e force field. The grid was then defined using the catalytic motif, i.e. residues 373-382 (whose side chains remained rigid), and the grid box size was 10 Å. It has been reported that MTM1 is more active towards lipids with acyl chains than only the headgroup [32,49], and hence the ligand used was PI3P with a short acyl chain (diC4; obtained from 1ZSQ MTMR2 lipid-bound crystal structure). A maximum of 10 poses was generated for each MTM1 protein and ligand pair. Following this, the free energy of binding for all the docked complexes predicted from XP glide docking was obtained using the Prime module from the Schrödinger suite, which calculates the free energy of binding or ΔG_binding_ in kcal/mol using the MM-GBSA method with default parameters [50]. The pose corresponding to the least ΔG_binding_ was selected as the top pose for each ligand-receptor complex and analysed using LigPlot+ [51,52]. The changes in PI3P-interacting residues across WT MTM1and the disease-causing mutants were traced. The interactions between the ligand backbone and residues within the catalytic pocket were represented as a scatter plot using the matplotlib module of Python3.5 (Teerna Bhattacharyya GitHub: https://github.com/teernab/MTM1_scripts/upload/main).

Although there exists information on a putative PI5P binding site which is implicated in aiding oligomerisation of MTM1 [9], the precise location of the binding site of PI3P in the GRAM domain of MTM1 is not known. Autodock 4.2.6 [53] was used to identify the putative PI3P binding site within the GRAM domain since XP glide docking cannot be performed without binding site information (‘blind docking’). With the minimised model of MTM1 as receptor and diC4-PI3P as the ligand, a round of rigid docking was carried out using Autodock 4.2.6, using 1000 steps of a genetic algorithm as the search method (default parameters), and 50 poses were generated (a higher number than that allowed for XP Glide docking since the aim of the blind docking exercise was to identify a putative ligand-binding site in the GRAM domain). The poses in which the ligand docked to the GRAM domain were analysed. In most top-ranking poses, the positively charged residues from the loops connecting the β5-β6 and β7-α1 were involved in interactions with PI3P. It had also been observed from DXMS studies on MTMR2 that these two loops are highly solvent accessible and, being on the same face as the MTMR2 active site, could participate in substrate binding [33]. The same could be extended for MTM1 owing to high sequence homology with MTMR2; the receptor grid was defined using the residues from these two loops in the GRAM domain of MTM1 and carried out XP glide docking of diC4-PI3P to the GRAM domain using the same parameters described above. Prime MM-GBSA method [50] was used to estimate the ΔG_binding_ of the PI3P-MTM1 docked complexes, and the docked pose with least ΔG_binding_ was analysed in LigPlot+ for the WT and the two GRAM domain mutants of MTM1.

### Mutagenesis, mammalian expression of MTM1 WT and measurement of the 3-phosphatase activity of MTM1 WT and mutant proteins using a mass-spectrometry based assay

The full-length sequence of human *MTM1* (NCBI Reference Sequence: NM_000252.1) cloned into pcDNA3.1(+) vector with an N-terminal FLAG tag was a kind gift from Dr Fred Robinson, OHSU, USA. Site-directed mutagenesis was carried out on the full-length *MTM1* as a template, using primers introducing single-nucleotide change, corresponding to each of the 14 point mutations across the MTM1 sequence. Primers of length varying between 39-54 nucleotides were designed to have a final Tm of about 78°C and GC content >30%. The PCR amplified products were purified and to confirm the desired point mutation and verified using Sanger sequencing. HEK293T cells grown at 37°C in 6-well plates in DMEM containing 10% FBS and PSG were transfected using Lipofectamine 3000 by the plasmids containing either WT or the mutant MTM1 constructs as per the vendor’s protocol. 36-48 hours post-transfection, the cells were harvested and lysed in phosphatase lysis buffer (20mM Tris-HCl pH 7.4, 150mM NaCl, 1% Triton X-100 (v/v) and protease inhibitor cocktail (Roche) as per Schaletzky et al., 2003 [9]. Total protein content in lysates was estimated using Bradford assay, and the protein expression of all constructs (WT MTM1 and mutants) for 20ug of total protein was checked using Western blot analysis. Nitrocellulose blots onto which protein bands from SDS-PAGE had been transferred were blocked using Blotto (0.1% Tween 20 in 1×TBS or TBST) and then incubated with anti-FLAG antibody (Thermo Fisher Scientific) and anti-GAPDH antibody (ImgeneX) overnight at 4°C and washed with 0.1% TBST before incubating with the secondary antibody (The Jackson Laboratory) at room temperature for 1.5 hours. The blots were developed using ECL Western blot medium (Thermo Fisher Scientific), and images were analysed using ImageJ or Fiji software (open source). The contrast of the images was enhanced only to visualise the protein bands of mutants showing poor expression. Quantification of these bands to understand the protein expression was carried out using raw images.

The lipid 3-phosphatase assay was performed as described in Ghosh et al., 2022 [54]. Briefly, 600 pmoles of 37:4 phosphoinositide 3-phosphate or PI3P (Avanti Polar lipids, LM-1900) were dried along with 20 uL of 0.5 M phosphatidyl-L-serine (Sigma, P5660) in vacuo. To 10 μg equivalent of protein (cell lysate) samples, phosphatase assay buffer wasadded. The reaction was incubated for 15 minutes at 37°C before adding 125 μLof 2.4 N HCl to quench the reaction. Clean phase separation was obtained through mixing and subsequent centrifugation at 1000 rcf for 5 minutes. The lower organic phase obtained was washed with 500 μL of lower phase wash solution and once again, the clean lower phase was obtained through mixing and centrifugation. Dried derivatised lipids were resuspended in methanol and assayed for odd-chain length acyl chain containing PI3P and PI using cells MRMs (995.5/613.5 Da for 37:4 PI3P and 809.5/613.5 Da for 37:4 PI). Controls used were blank (only solvent), untransfected HEK293T cell lysate (UTC) and heat-killed WT MTM1 (HK; WT MTM1 cell lysate heat denatured at 95°C). The run conditions used were as per Ghosh et al., 2019 [55] and Ghosh et al., 2022 [54]. The lipid 3-phosphatase activity of the proteins was measured as the ratio of PI and PI3P released post-reaction, indicating the extent of dephosphorylation of PI3P by the WT and mutant MTM1 proteins. Data were statistically analysed using MS-Excel (Office 2016), and one-way ANOVA with posthoc Tukey’s multiple comparison test was used for significance testing.

## Acknowledgements

The authors would like to thank NCBS (TIFR) for infrastructure and other facilities. RS acknowledges support from Institute of Bioinformatics and Applied Biotechnology through her Kiran Mazumdar Shaw Computational Biology Chair position.

